# Uncovering variation in social tolerance among wild vervet monkeys through a novel co-feeding paradigm

**DOI:** 10.1101/2025.08.03.668336

**Authors:** Fabio Opreni, Erica van de Waal, Rachel Harrison

## Abstract

Social tolerance, defined as the probability of individuals maintaining proximity with minimal aggression, is essential for the functioning of social groups. This study investigates social tolerance in wild vervet monkeys (*Chlorocebus pygerythrus*), with a focus on co-feeding tolerance. Previous research has highlighted that social tolerance varies both between and within species, influenced by stable group-specific and temporally variable factors. However, it remains unclear how these differences manifest in wild primates, particularly among vervet monkeys. To address this, we employed a novel experimental co-feeding paradigm adapted for field conditions, presenting a fake grass carpet with a fixed density of corn and a plot area proportional to group size to measure group-level co-feeding tolerance. The study was conducted with two habituated groups of wild vervet monkeys at the iNkawu Vervet Project in South Africa. Results revealed significant group differences in co-feeding tolerance despite similarities in the groups’ demographics and environments. The presence of mothers with infants was associated with higher levels of co-feeding tolerance, though this effect differed by group. These findings highlight the need for further research to elucidate the factors driving group-specific social tolerance levels and the mechanisms behind these differences.

## Introduction

Social tolerance is the probability of individuals being in proximity to each other with little or no aggression (1). It is a key feature needed in social groups, likely benefitting individuals through protection against predators, reduction of uncertainty to better cope with unpredictable environments, and arguably better foraging efficiency (2). It is a fundamental principle allowing animal societies to function and potentially makes it possible to develop and maintain complex cooperative behaviours (3). Successful group living requires individuals to manage their behaviour in ways that avoid hindering other members and generating conflict. Indeed, for species depending on group cohesion, the level at which individuals achieve social tolerance is linked to their survival and reproduction, hence their individual fitness(4) Indeed, much evidence underscores the importance of social tolerance in social species across the entire animal kingdom for the success and cohesion of a given group (5,6).

For individuals to function effectively as a collective, costs such as within-group competition must be minimized. This enables close interactions and the avoidance of conflict (7,8). Social tolerance is often considered a result of successfully achieving this balance (1). However, in the non-human primate literature, there are a variety of potential methods and approaches regarding how to assess social tolerance. It has been measured, for instance, using specific tolerant behaviours such as food sharing, a voluntary transfer of food from one individual to another (9,10), or via experimental methods such as co-feeding paradigms, an experimental feeding in a restricted area, forcing individuals to be close to each other, to measure co-feeding tolerance as a proxy for social tolerance (11,12). It has also been measured using the structural specifics of a group, such as the covariation of affiliative and aggressive interactions, as well as dominance relationships (13–15) underlining the importance of distinguishing between social tolerance as a structural construct and as a behavioural construct. Structural social tolerance is measured by specific behaviours or traits (e.g., grooming, aggressions) that are not explicitly tolerant, while behavioural social tolerance is measured by inherently socially tolerant behaviours (e.g., food sharing, co-feeding), described as peaceful, non-agonistic interactions between individuals (13). Behavioural social tolerance is considered a distinct concept, but it is enabled by structural social tolerance (13).

According to the Relational Model, behavioural social tolerance is one of three outcomes of social conflict along with conflict escalation and conflict avoidance, attesting to its importance in the frame of competition and conflict resolution for group success (16). The co-feeding paradigm simulates resource competition and the potential for social conflict over a resource, revealing how much of the outcome, according to the Relational Model, is social tolerance, and is a reliable and frequently used method, representing a good measure of the manifestation of behavioural social tolerance, allowing both group-level and dyadic-level analysis (13). Measuring group-level co-feeding tolerance as a proportion of the group that co-feed compares the social tolerance outcome (co-feeding individuals) with what is approximated as conflict avoidance (individuals from the group not coming to co-feed) (17). Co-feeding tolerance is arguably therefore a good approximation of social tolerance.

The levels at which competition and cooperation act on social tolerance have long been studied at the species level. This has been broadly studied between related primate species, for instance, between Sumatran (Pongo abelii) and Bornean orangutans *(Pongo pygmaeus)* (18), red-fronted lemurs *(Eulemur rufifrons)* and ring-tailed lemurs *(Lemur catta)* (19), and bonobos *(Pan paniscus)* and chimpanzees *(Pan troglodytes)* (20). Macaque species have been categorized according to their tolerance level (21). However, social tolerance does not only vary between species; evidence strongly suggests that, even if there is a species-specific component, social tolerance varies between groups within a species (12). However, most of our understanding to date regarding social tolerance variation within species comes from captive settings, where constant food availability and human habituation are known to alter social structures (19,22). Different studies have emphasized the critical role of ecological validity in primate cognition research, highlighting that natural environments offer more complex and diverse social interactions (23,24). Therefore, assessing within-species social tolerance variation in the wild would be a valuable contribution to the literature, especially when identifying selective pressures (25).

In the context of co-feeding tolerance, the rates of co-feeding and proximity between individuals can be modified by potential dominant individuals if the food source can be monopolized. It is thus important to apply a method that allows an opportunity for every individual in a group to participate (26). Group size is indeed expected to create more foraging competition, but in primates, it is not necessarily associated with different aspects of group sociality or a difference in social tolerance (27,28). However, to our knowledge, co-feeding paradigms controlling for group size have only been applied in captive or semi captive (12) conditions but never in the wild.

Previous research suggests a relationship between group composition, particularly the presence of mothers with infants and young females, and levels of social tolerance (12). While these demographic factors explain some variance in observed social tolerance, group-specific differences persist even under similar ecological conditions (12,26). There is substantial evidence that primate groups exhibit culturally distinct behaviours, including tool use, hunting strategies, and other socially transmitted practices (29). Such cultural differences include social behaviours, with variations in aggression and affiliation patterns observed between groups (30,31). These group-unique social styles may reflect underlying differences in structural social tolerance. Indeed group specific dynamics can be shaped by the behaviours of individual members and can be explained by characteristics such as the influence of dominant individuals, who play a crucial role in behavioural transmission within the group (32), and success-biased learning, where individuals preferentially adopt behaviours from successful group members (33).

There is evidence for variation in social behaviour between groups of wild vervet monkeys. Kerjean and colleagues (34) using nine years of observational data from the iNkawu Vervet Project, South Africa (henceforth: IVP), highlighted consistent differences in social behaviour among three neighbouring groups. Earlier findings had indicated differences in grooming networks (35) and other social network parameters between these groups (36), with both studies identifying Ankhase (AK) as exhibiting higher structural social tolerance than the other groups. Kerjean and colleagues similarly found that AK’s sociality is characterized by a higher propensity for affiliative behaviours, greater grooming reciprocity, and interactions across wider rank distances, traits that distinguish AK as an outlier in structural social tolerance (34–36).

However, there is no experimental evidence of group differences in co-feeding tolerance using a paradigm controlling for group size in this population of wild vervet monkeys (or, to our knowledge, in any population of wild primates). Moreover, there is no evidence regarding how differences in group composition, such as the presence of mothers with infants, might be associated with co-feeding tolerance variation in this species. Observations of field experiments conducted in two wild groups of vervet monkeys sharing similar ecological conditions at IVP will provide an ecologically relevant setting to study co-feeding tolerance differences and covarying factors.

To further investigate whether group differences in social dynamics are driven by demographic factors, we included KB, a group not studied by Kerjean and colleagues. KB shares key demographic characteristics with AK, including a similar sex ratio, group size, and number of new-born infants, all of which are group composition variables known to affect group sociality (12,26,37). Comparing these two groups allows us to control for demographic variation and potentially identify other factors contributing to differences in co-feeding tolerance.

Based on previous literature, group-level co-feeding tolerance is measured according to the proportion of the group co-feeding across the experiment time, providing an estimation of behavioural social tolerance (12,13,26). According to the Relational Model (16) we will gain insight into how much a potential for conflict leads to behavioural social tolerance (co-feeding individuals) compared to what would be estimated as conflict avoidance (individuals not participating) and how much it leads to social conflict, based on co-feeding related aggressions (17).

We hypothesize that the AK group will exhibit higher co-feeding tolerance, measured by the proportion of group members feeding simultaneously, compared to the other group. This prediction is informed by previous findings indicating that AK individuals tend to display higher structural social tolerance (34–36), and aligns with broader evidence that social tolerance varies between primate groups even under similar ecological conditions (12,26). Secondly, we hypothesize that higher group-level tolerance in AK will correspond to lower rates of aggression during co-feeding experiments. Finally, we predict that the presence of mothers carrying infants, a key demographic factor, will positively influence co-feeding tolerance in both groups, consistent with findings in chimpanzees (12).

## Methods

### Ethics

This research adhered to the Association for the Study of Animal Behaviour Guidelines for the Use of Animals in Research, and was approved by the relevant local authority, Ezemvelo KZN Wildlife, South Africa.

### Study site and subjects

The study was conducted at the iNkawu Vervet Project (IVP) research organization and field site, situated in Mawana Game Reserve in KwaZulu-Natal, South Africa (S 28° 00.327; E 031° 12.348). Two neighboring groups of wild vervet monkeys habituated to human observers (Ankhase (AK) and Kubu (KB)) were used for data collection. These groups varied in the number of individuals from September 2023 to March 2024 (see Figure 1). The maximum group sizes were 20 for AK and 22 for KB. The variation in group size during the study period was due to deaths, male dispersals, and immigrations. A variation in the observed group size could also be due to the failure to spot some individuals on the day of the experiment.

**Figure 1.**
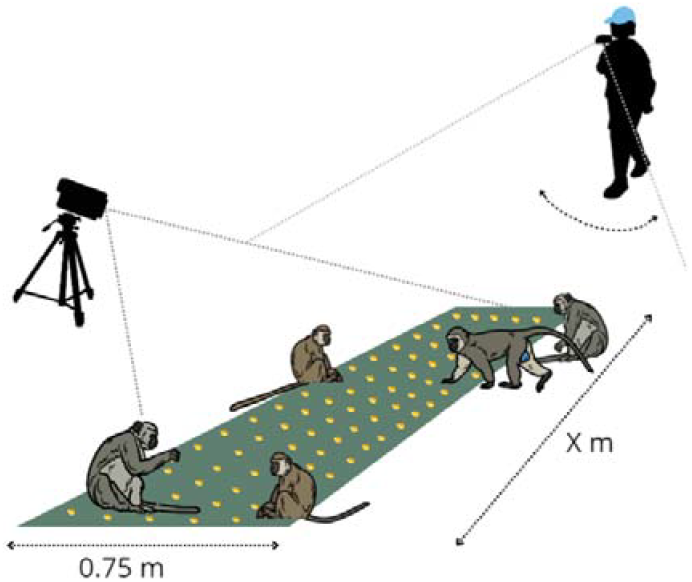
Experiment set-up consisted of a fake grass feeding plot with a fixed density of food. It was 0.75 meters wide and 2.7 meters long for AK & KB.

The birth season lasted approximately from October 2023 to December 2023, during which mothers carrying infants could be observed. New-born infants were not considered in the group size; however, mothers carrying their infants were recorded when coming to co-feed.

### Experimental set-up & data collection

The experimental setup was inspired by experiments conducted in semi-captive conditions on chimpanzees and bonobos (see: Staes et al., 2022; DeTroy et al., 2021) and adapted for vervet monkeys in the wild. The experimental setup consisted of a food source plot of a specific length according to the group size that was presented to the monkeys. The food source plot consisted of a 0.75 m wide fake grass carpet, with a length dependent on the group size, set at 0.123 m per individual. The length of the fake grass carpet for both groups was 2.7m (Table 1).

**Table 1.** Group characteristics concerning experimental setup and data collection. Each experiment ranged from 1 up to 8 scans. The feeding plot length was assessed proportionally to the group size.

|  | Ankhase (AK) | Kubu (KB) |
| --- | --- | --- |
| Number of experiments | 13 | 15 |
| Total number of scans | 62 | 67 |
| Group size range across time | 18-20 | 20-22 |
| Feeding plot length | 2.7 m | 2.7 m |

The food reward offered consisted of overnight-soaked corn, which the monkeys are habituated to (see van de Waal, et al. 2013). The density of corn on the carpet was set at 12 pieces per individual.

The fake grass carpet was unrolled near the sleeping site early in the morning when the group was gathered there. The plot was placed according to the vegetation, ensuring that the carpet lay flat on the ground but was close enough to thicker vegetation or trees to provide a safe place for the monkeys to approach the food. In the summer period, which is the rainy season, the denser vegetation often required the use of a machete to clear an area for placing the fake grass carpet. The rainy season started in November 2023 and lasted until the end of the data collection. Two pilot trials were initially conducted with both groups to habituate the monkeys to the experimental setup, with no data collected during these trials.

Each session was recorded by one camera on a tripod and another mobile camera carried by a person to provide different angles of recording. At least one observer, qualified to recognize the monkeys, narrated the names of the individuals co-feeding on the plot from left to right, as well as the number of visible aggressions every 30 seconds. The monkey identification test for a specific group is standardized for all IVP field site researchers to ensure accurate identification of individuals. The observer(s) were positioned on the opposite side from where the monkeys were, to provide a safe approach (see Figure 1 for the setup visualization). The total group size was assessed according to the observation form from the day before. Experiments were aborted if there was a temporary group split, an alarm event, or a group encounter registered by the field site researchers.

To attract the monkeys to the food, researchers used a specific call that the monkeys have previously learnt to associate with food. A session began when the first individual took a piece of food and ended either when all the corn was consumed or when the monkeys left the feeding area. The number of individuals co-feeding was recorded at 30-second intervals, representing one scan, resulting in up to 8 scans per session. The first scan was taken 30 seconds after the beginning of the session. The fake grass carpet was removed once the monkeys lost interest in it and were no longer in sight.

In total, 13 experiments were conducted with AK and 15 with KB; all were evenly distributed across September 2023 to March 2024.

### Video coding

Video coding was conducted using VLC player and transcribed into Excel spreadsheets.

Group co-feeding tolerance was defined as the number of individuals per scan within arm’s length of the food source (and hence capable of reaching the corn) over the total group size accounted for the day before (therefore providing a proportion of the group present).

All co-feeding monkeys were identified and counted every 30 seconds (corresponding to one scan) resulting in a list of all monkeys feeding during a given scan. The individual recognition, on site, and during the video coding was performed with the competence of an IVP identification test for the group. The number of aggression events was coded from each scan either from the speaking of a field site researcher or if seen on the video, according to the list of aggressive behaviours listed in Table 2.

**Table 2.**
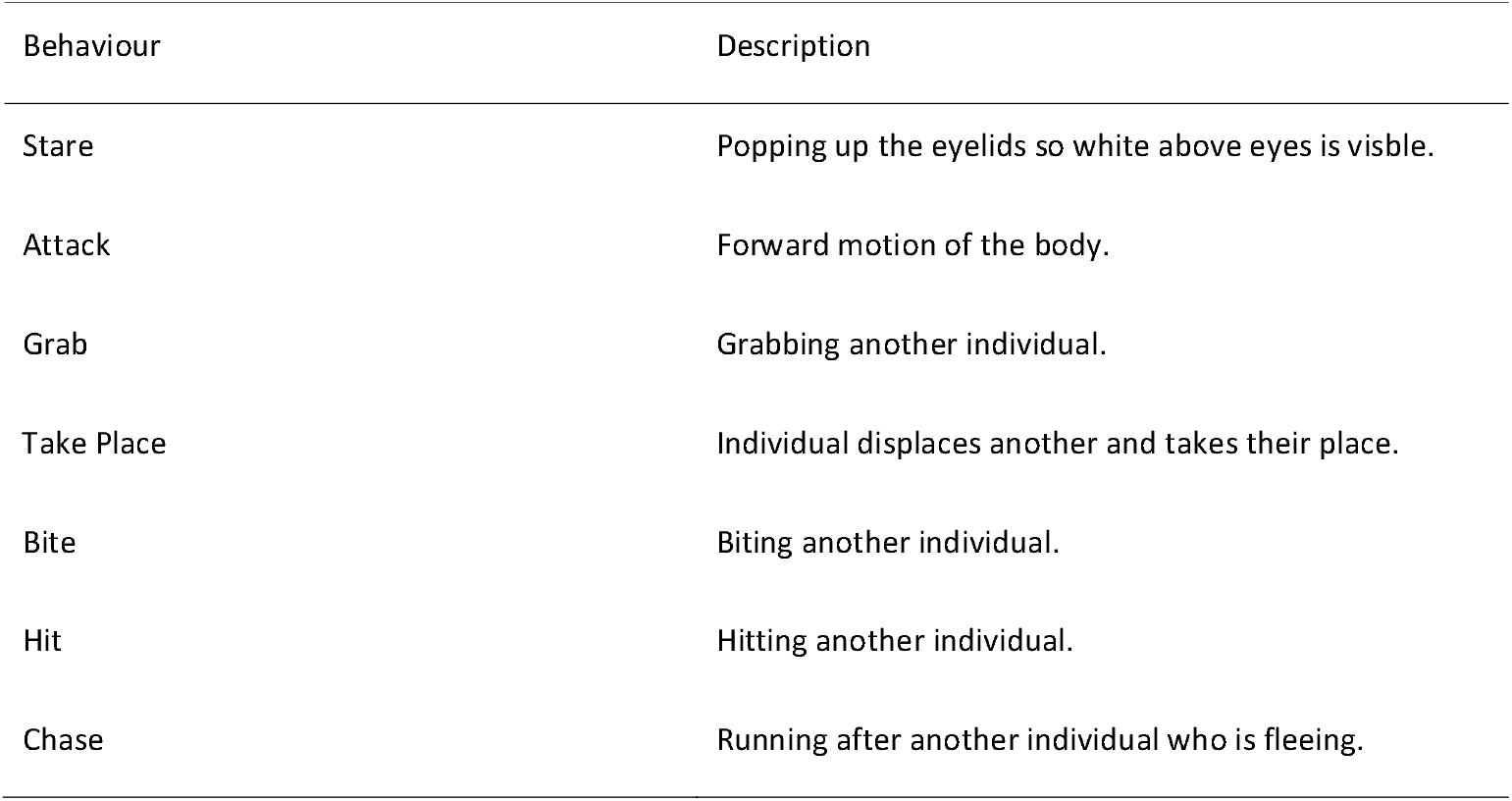
List of aggressive behaviours considered in the number of per scan aggressive events.

### Statistical analyses

Statistical analyses were conducted using R (39) and Rstudio (40). All graphs were created using the ggplot2 package (41). Generalised linear mixed models (GLMMs) were used to analyse the proportion of each group co-feeding during experimental sessions and the frequency of aggression during experimental sessions The proportion of each group co-feeding at each scan during each experimental session was assessed using a binomial GLMM with logit link function (function ‘glmer’ in the R package lme4; (42). Group (AK vs. KB) and Scan (z-transformed to have a mean of 0 and standard deviation of 1) were included as main effects, with a random slope of Scan by Experimental Session nested within Group.

The count of aggressive interactions during each scan was assessed using a poisson GLMM with a log link function. Group (AK vs. KB) and Scan (z-transformed to have a mean of 0 and standard deviation of 1) were included as main effects, with a random effect of Experimental Session nested within Group.

The effect of the presence of mothers carrying infants upon the proportion of each group co-feeding at each scan during each experimental session was assessed using a binomial GLMM with logit link function (function ‘glmer’ in the R package lme4; (42) Bates et al., 2014). Group (AK vs. KB), Presence of mothers with infants (Present vs. Absent) and Scan (z-transformed to have a mean of 0 and standard deviation of 1) were included as main effects, with an interaction between Group and Presence of mothers, and the model included a random slope of Scan by Experimental Session.

All GLMMs were assessed using full – null model comparisons, and multicollinearity was assessed using variance inflation factors (function ‘vif’ in the R package car; (43). Variance inflation factors below five were considered acceptable and the highest variance inflation factor observed was 2.7. For the poisson GLMM analysing counts of aggression, overdispersion and zero inflation were assessed using the ‘testDispersion’ and ‘testZeroinflation’ functions in the R package DHARMa (44) and no issues were identified.

## Results

### Comparisons of co-feeding proportions

The full model was a significantly better fit to the data than a null model containing only the random effects structure (^2^ =13.62, *p* = 0.001). A binomial GLMM indicated that KB had a significantly lower proportion of the group co-feeding across the experimental sessions than AK (= −0.25, *p* = 0.018, see Figure 2). There was also a negative effect of scan in both groups (= −0.22, *p* = 0.004), indicating that monkeys tended to co-feed at lower proportions as experiments ran on (likely due to resource depletion).

**Figure 2.**
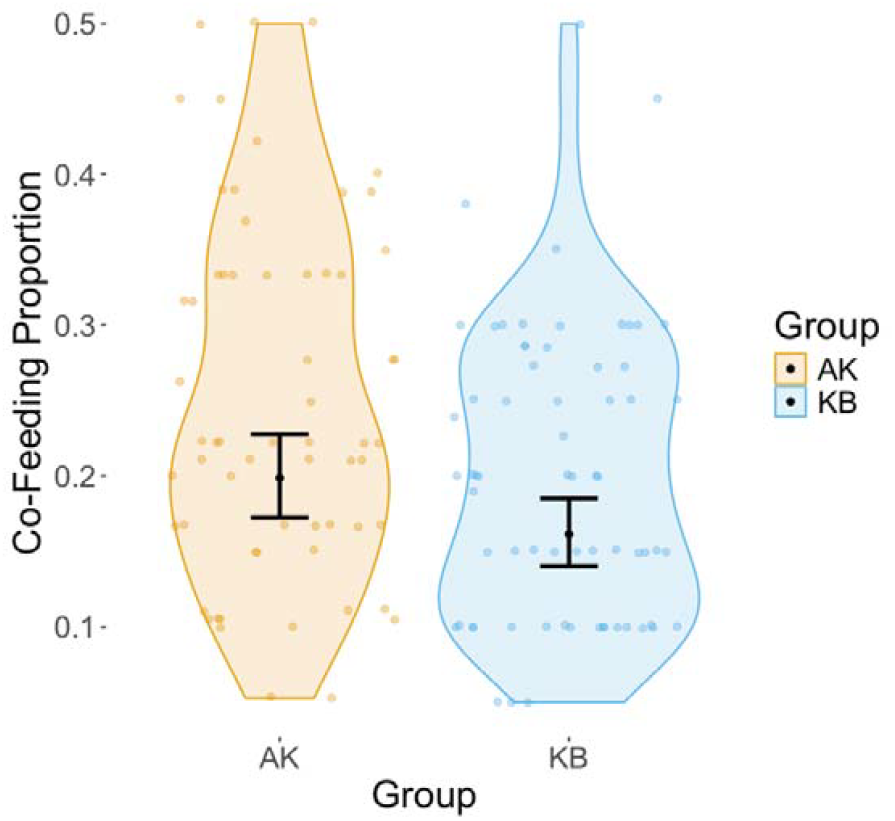
Violin plots show the observed proportion of each group co-feeding at each scan across all experimental sessions. Points show the observed proportion at each scan. Black points show the predicted proportions from the GLMM with error bars showing the 95% confidence interval around these predictions.

### Comparison of aggression during experiments

The full model was a significantly better fit to the data than a null model containing only the random effects structure (^2^ = 8.15 p = 0.043). The interaction between Scan and Group was also close to significance (= 0.53, p = 0.088), as shown in Figure 3, with aggression increasing over the course of each experiment in KB but remaining stable in AK. At the mean value of Scan, AK and KB did not differ significantly in their counts of aggression during experiments, though this tended towards significance with KB having slightly higher frequencies of aggression (1.00, p = 0.065). A model without this interaction term was not a significantly better fit to the data than a null model containing only the random effects structure (^2^ = 5.17, p = 0.08), and found no significant main effect of = Group upon frequency of aggression (= 0.96, p = 0.07).

**Figure 3.**
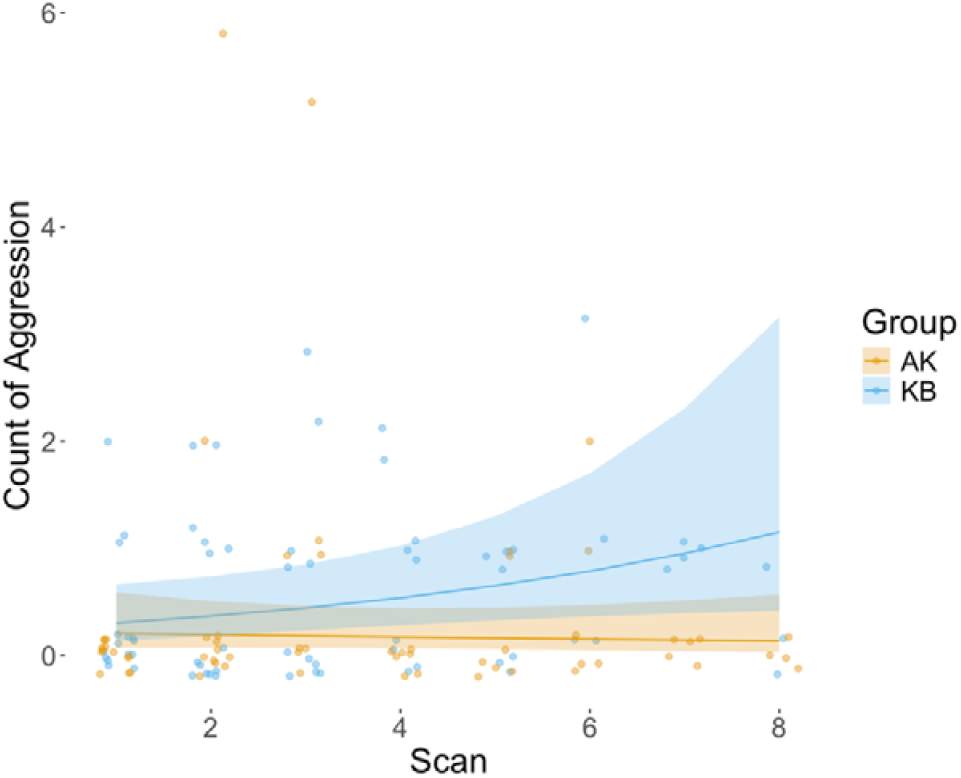
Points show the observed counts of aggressive interactions at each Scan across all Experimental Sessions. Solid lines show the predicted values from the model, with shaded ribbons showing the 95% confidence interval around these predictions.

### Impact of mothers carrying infants

There was a significant interaction between Group and the effect of mothers carrying infants, such that the presence of mothers with infants was linked to higher co-feeding proportions in KB (= 0.60, p = 0.014), but had no effect in AK (= 0.14, p = 0.45, see figure 4).

**Figure 4.**
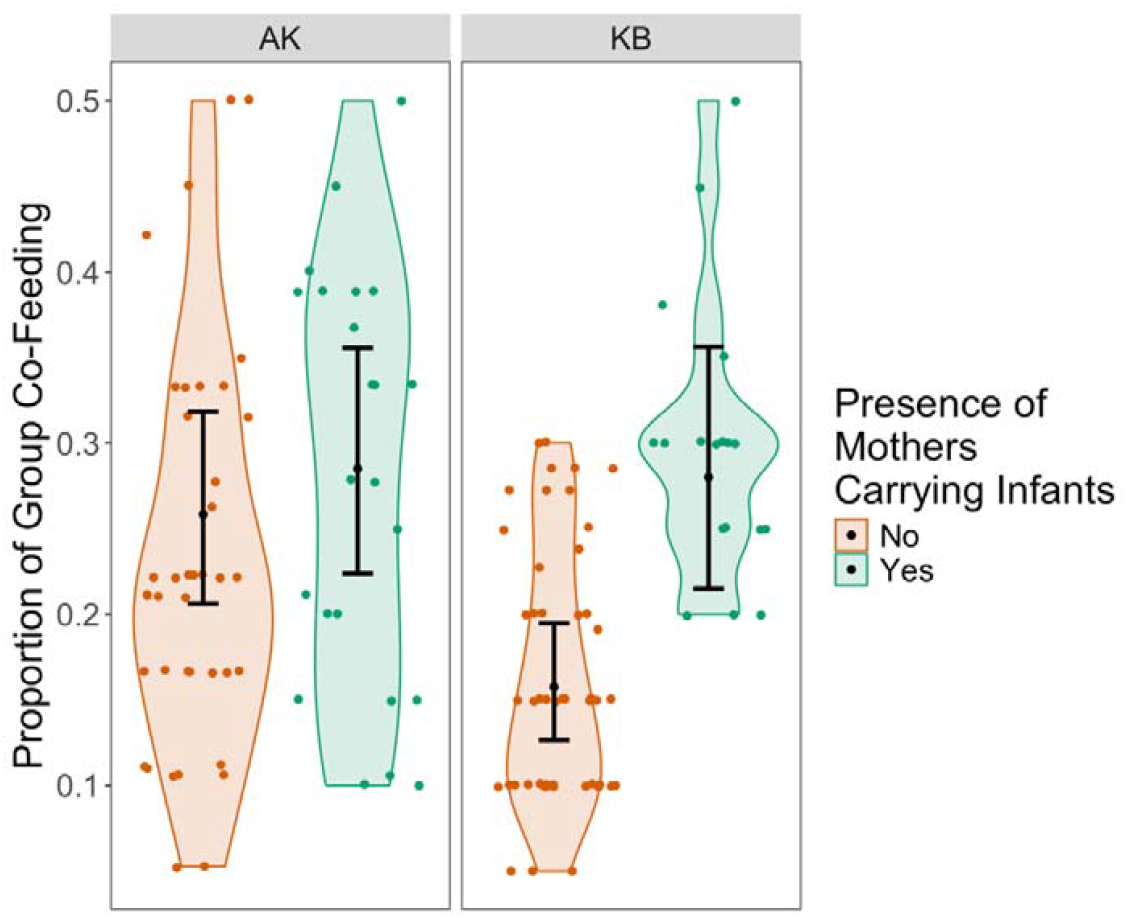
Violin plots show the observed proportion of each group co-feeding at each scan across all experimental sessions when mothers carrying infants were present versus absent. Points show the observed proportion at each scan. Black points show the predicted proportions from the GLMM with error bars showing the 95% confidence interval around these predictions.

## Discussion

This study provides insights into variation in social tolerance among wild vervet monkeys using a novel experimental co-feeding paradigm, highlighting the flexible nature of co-feeding tolerance within a species. Notably, we found significant group differences in co-feeding tolerance: AK consistently exhibited higher levels of co-feeding tolerance compared to KB, despite the two groups sharing similar demographic characteristics. Group composition, specifically the presence of mothers with their infants on the feeding plot, was associated with higher levels of co-feeding tolerance in KB.

While this study provides the first experimental evidence of group-level differences in social tolerance in wild vervet monkeys, there are important limitations to consider. First, social tolerance was assessed in the context of co-feeding tolerance, which may limit generalizations and comparisons with other research that uses different measures Co-feeding tolerance is highly context-specific and social tolerance might not function the same wa in other contexts. Secondly, group-level co-feeding tolerance was assessed using the proportion of co-feeding individuals within the group, which can be biased by demographic changes. This measure also relies on the assumption that all group members are present and have the same opportunity to co-feed. The accuracy of this measure depends on researchers having correctly identified all individuals the day before and on all group members sleeping in the same location.

Demographic changes are associated with temporally variable behavioural traits, including co-feeding tolerance. This suggests that shifts in group characteristics can influence co-feeding tolerance changes over time (12,45). Indeed, longitudinal variations in co-feeding tolerance were found to be non-random in chimpanzees, particularly with the presence of mothers with young infants, which was associated with a decrease in co-feeding tolerance (12). In contrast, our results from AK and KB showed a different pattern: we found that the presence of mothers carrying infants was associated with significantly higher co-feeding tolerance in KB, but not in AK.

One possible explanation for this difference is that AK may have already reached a consistently high level of co-feeding tolerance, leaving little room for further measurable increases. Although the carpets were scaled to group size, it is worth questioning whether the setup truly allowed all individuals to feed at once. Even with proportional scaling, individual spacing needs or social dynamics may still have limited full group co-feeding. As a result, co-feeding tolerance in AK may have reached a threshold beyond which further increases were not possible.

This result contrasts with the trend found in chimpanzees by DeTroy et al. (2021), who suggested that mothers with infants exhibit increased vigilance due to the vulnerability of their young and therefore refrain from interacting in proximity to others (12,46). In chimpanzees, young infants face a high risk of infanticide from conspecifics, including both males and females, whether within or outside the group (47,48). In contrast, infanticide in vervet monkeys is rare and typically involves newly immigrated males (49), and it has never been observed in the IVP population during 14 years of observation (E. van de Waal, *personal communication*). This lower threat level may account for the differing results. Moreover, vervet monkeys show high curiosity toward new-borns, and mothers carrying infants often receive increased affiliative attention (50,51). This likely leads to increased proximity between mothers and other group members, potentially contributing to higher group-level social tolerance, as reflected in the significant effect found in KB.

Our results also suggest that group demographics alone do not account for the differences in co-feeding tolerance. For example, KB and AK shared similar demographics (i.e., similar group sizes with minimal variation and similar sex ratios), yet their co-feeding tolerance differed, with AK showing significantly higher values. Potential effects of differing local ecological environments cannot be entirely excluded, as the two groups inhabit different territories with potentially varying resource availability (52). However, given that the global habitat has similar environmental characteristics, it is unlikely that habitat heterogeneity alone explains these neighbouring group differences. Between-group aggression levels seem to support the co-feeding tolerance results specific to each group, according to the Relational Model, which suggests that lower co-feeding tolerance is associated with higher aggression levels. While not statistically significant, aggression patterns diverged between AK and KB over the course of each experiment. In AK, aggression remained consistently low, whereas in KB it increased over time. This rise might reflect growing competition as food on the carpet became scarcer, suggesting group-specific differences in how competition is socially managed.

Genetic factors are unlikely to influence group differentiation, as these neighbouring groups experience regular gene flow due to dispersing males (38). Group-specific characteristics can therefore be expected, and literature refers to these as “collective personalities” (53,54). Such group differentiations, where demographic, genetic, and ecological factors do not account for all variability, can be attributed to learned behavioural styles (55). Previous work has consistently shown that AK exhibits higher affiliative tendencies, broader social engagement across rank distances, greater grooming reciprocity, and elevated co-feeding tolerance compared to other groups (34–36). These repeated patterns across independent studies suggest that AK stands out as an outlier in terms of structural social tolerance and hint at the possibility of a socially learned group behavioural style, what may be considered a social tradition. However, the mechanisms underlying such traditions remain to be formally investigated.

In AK particularly, the consistently high social tolerance results compared to other groups lasting over 10 years might also be due to a social tradition lasting multiple generations. Indeed, only one individual from the start of IVP remains as an adult and never participated in my experiments, so the consistent presence of influential individuals cannot be responsible for the observed pattern. Social traditions lasting multiple generations are not uncommon in non-human primates. There is for instance evidence regarding tool use (56), communication (57) and specific affiliative behaviours (58).

However, further research is needed to disentangle the extent to which variation in group differences arises from temporal factors versus group-specific factors. Additionally, there is a lack of evidence on the extent to which learned behavioural styles contribute to collective personalities.

### Conclusion & perspectives

This study highlights the within-species variation in co-feeding tolerance among wild vervet monkeys and emphasizes the need to distinguish between the contributions of temporal factors and stable group-specific factors to this variation. Factors such as seasonality and group composition, particularly the presence of mothers with infants, likely influence temporal variation in co-feeding tolerance. However, further research is needed to identify additional aspects affecting this variation. For example, components of group composition like the presence of young females have been linked to social tolerance (12), and sex ratio has also been found to influence social tolerance levels (59). A long-term analysis would help disentangle these factors.

Further research should also explore the components leading to group-specific behavioural styles or “collective personalities” (60) to better understand the reasons behind stable group differences.

## Acknowledgements

We thank all members of the IVP field team for their help with data collection. Special thanks to Michael Henshall, IVP field site manager, for his assistance in scheduling the experiments.

